# Measuring and Modeling Macrophage Growth using a Lab-on-CMOS Capacitance Sensing Microsystem

**DOI:** 10.1101/2023.01.27.525955

**Authors:** Kyle Smith, Ching-Yi Lin, Yann Gilpin, Elizabeth Wayne, Marc Dandin

## Abstract

We report on the use of a lab-on-CMOS biosensor platform for quantitatively tracking the growth of RAW 264.7 murine Balb/c macrophages. We show that macrophage growth over a wide sensing area correlates linearly with an average capacitance growth factor resulting from capacitance measurements at a plurality of electrodes dispersed in the sensing area. We further show a temporal model that captures the cell evolution in the area of interest over long periods (e.g., 30 hours). The model links the cell numbers and the average capacitance growth factor associated with the sensing area to describe the observed growth kinetics.

## 1 Introduction

Macrophages are innate immune cells that specialized inflammatory functions related to damage detection, pathogen recognition, clearance, and wound healing (Wynn, Chawla and Pollard, 2013). To perform these heterogeneous functions, macrophages undergo polarization, whereby cells produce a specialized phenotype in response to environmental stimuli. Tissue-resident macrophages become polarized during the progression of infectious disease and chronic diseases like cancer, neurodegeneration, and autoimmunity (Williams *et al*., 2018; Ma *et al*., 2019; Reis-Sobreiro *et al*., 2021). As such, drug targeting for the purpose of re-polarizing macrophages is a major basic and translational research area. Building a tool to conduct a real-time investigation of macrophage polarization response to biomolecular cues within their microenvironment is of paramount importance for the development of novel therapies.

Current macrophage polarization characterization methods are accurate but result in a loss of temporal resolution. For example, flow cytometry, qPCR, proteomics, and transcriptomics techniques measure genetic and protein content but require tissue homogenization or fixation (Derlindati *et al*., 2015; Tarique *et al*., 2015; He *et al*., 2021). This limits the opportunities for repeated measurement. *In situ* fluorescent or colorimetric labeling of cells can be used to assess macrophage polarization, but these dyes can potentially alter macrophage functions. Further, staining is usually performed as an endpoint assay, making it difficult to track cells in real time over extended periods (Saylor *et al*., 2018). Furthermore, these techniques require skilled laboratory technicians and significant hardware overhead for proper execution. These two factors render them expensive and difficult to deploy in resource-limited settings.

The use of electronic microsystems to monitor biological processes helps mediate these issues. Biosensors, as they are termed, are electronic microsystems that provide non-destructive and label-free detection of cell health in response to environmental changes (Naresh and Lee, 2021). In comparison to the state-of-the-art, they are easy to use, and they are relatively low in cost.

In this paper, we describe the use of a lab-on-CMOS biosensor for studying macrophage dynamics in their growth phase. A lab-on-CMOS device is a platform that integrates lab-on-a-chip technology with complementary metal-oxide semiconductor (CMOS) chips for biosensing (Roussel *et al*., 2006; Christen and Andreou, 2007; Sawan, Miled and Ghafar-Zadeh, 2010; Ghallab and Ismail, 2014; Yin, Li and Mason, 2016; Mason and Wan, 2017; Li, Yin and Mason, 2018; Lin *et al*., 2022). The chips can feature circuits that are configured to transduce biophysical and biochemical events to the electrical domain and signal processing hardware for measurement, conditioning, and reporting. The biological species under test, in our case the macrophages and their liquid media, are applied directly on the chip’s surface where a plurality of sensors are disposed. Other parts of the chip are isolated from the cell media using a custom-designed package configured to preserve the chip’s electrical integrity while it is operating in a wet environment.

Our chip’s transduction mechanism is based on interfacial capacitance sensing, which allows the tracking of cell viability, proliferation, and morphology changes (Forouhi, Dehghani and Ghafar-Zadeh, 2018, 2019). The principle of operation of interfacial capacitance sensing is like that of electrical cell-substrate impedance sensing (ECIS) (Susloparova *et al*., 2013, 2015; Hu, Arcadia and Rosenstein, 2021). ECIS measures changes in impedance at a sensing electrode as a function of frequency and time, whereas interfacial capacitance biosensing tracks changes in capacitance at the sensing electrode as a function of time (Hedayatipour, Aslanzadeh and McFarlane, 2019).

While both methods can yield the same information about a cell culture overlying a set of sensing electrodes, capacitance sensing circuit architectures are simpler to implement because the complex current sourcing and wide band frequency scanning circuitry typically used in ECIS schemes are not needed to sense changes in capacitance (Hedayatipour, Aslanzadeh and McFarlane, 2019). Rather, simpler topologies can be employed to convert the sensed capacitance into a voltage, a current, a frequency, or a pulse width (Ferlito *et al*., 2020).

Capacitance sensing bioelectronics is well established. For example, it is used in the biopharmaceutical industry to improve scale-up cell culture processes (Konakovsky *et al*., 2015; Reinecke *et al*., 2017; Metze *et al*., 2020). This includes using capacitance-sensing probes to study and detect biomass and viable cell concentrations in suspensions and to determine how close to confluency a batch is. However, these use cases typically do not include adherent cell lines and much less the ability to detect cell counts from capacitance measurements.

Lab-on-CMOS capacitance sensors have also been demonstrated previously (Forouhi, Dehghani and Ghafar-Zadeh, 2019). They are advantageous because they allow smaller sample volumes to be analyzed. And, a plurality of analysis functions and signal processing can be integrated on the CMOS chip, which makes these microsystems ideal for point-of-care applications. Their use has been shown in drug cytotoxicity assays (Nabovati *et al*., 2019), potency assays for chemotherapeutic agents (B. Senevirathna, Lu, Dandin, *et al*., 2019), viral infection assays (Abdelhamid *et al*., 2022), oral cell analysis (Osouli Tabrizi *et al*., 2022), and nanoparticle-mediated activation of neutrophils (Bunnfors *et al*., 2020).

Furthermore, the cell coverage of an electrode and its relationship with the elicited capacitance change at the electrode has been studied in an attempt to quantify the relationship between measured capacitance and cell density in a capacitance sensing lab-on-CMOS device. For example, Senevirathna *et al*. showed that the measured capacitance at an electrode was correlated with the cell coverage at that electrode (B. P. Senevirathna *et al*., 2019). Further, Renegar *et al*. developed a framework based on deep neural network image segmentation techniques to study the correlation between the measured capacitance and the coverage of single cells at an electrode (Renegar, Noyan and Abshire, 2022).

These works offer compelling evidence that biophysical phenomena (e.g., cell spreading and coverage) modulate interfacial capacitance. In this paper, we present a methodology for deriving a temporal model that can predict cell growth over a wide area based on an average capacitance inferred from measurements originating from a plurality of electrodes disposed inside the area. This is unlike the aforementioned studies, which focused on local effects at the electrode sites. In contrast, we show that in sustained growth conditions, the set of sensing electrodes can be considered as one electrode registering an average capacitance growth factor over the course of the culture. We further show that the average capacitance growth factor and the cell counts registered over the area can be linked via a temporal model that tracks the evolution of the cell numbers inside the area of interest. This model and the data analysis techniques featured in this paper lay the groundwork for a generalized framework for gaining insights from capacitance sensors using multiple electrodes over a wide sensing area.

## 2 Materials and Methods

### 2.1 Lab-on-CMOS Biosensing Platform

The biosensing platform used in this study included a microsystem configured for measuring cell proliferation and migration (Senevirathna *et al*., 2016; B. P. Senevirathna *et al*., 2019; B. Senevirathna, Lu, Dandin, *et al*., 2019; Gilpin *et al*., 2022). At its core, the microsystem included an application-specific integrated circuit (ASIC) fabricated in a 0.35 μm CMOS technology. The ASIC chip included 16 capacitance-to-frequency (CTF) sensors structured in a 4 × 4 array of integrated biosensor pixels with a spatial pitch of 196 × 186 μm. Each pixel included an interdigitated electrode structure with a sensing area of 30 × 30 μm^2^ and dedicated circuitry for transducing cell adsorption, movement, or life cycle events (e.g., mitosis) into an electrical signal. The electrodes were isolated from the cell media using the CMOS process’ native passivation layer. The sensors measure the capacitance at the cell interface, and they generate a digital signal that is outputted to an off-chip microcontroller via an on-chip I^2^C interface. The sensors’ principle of operation is described below, referring to Figure 1(A, B, and C).

**Figure 1.**
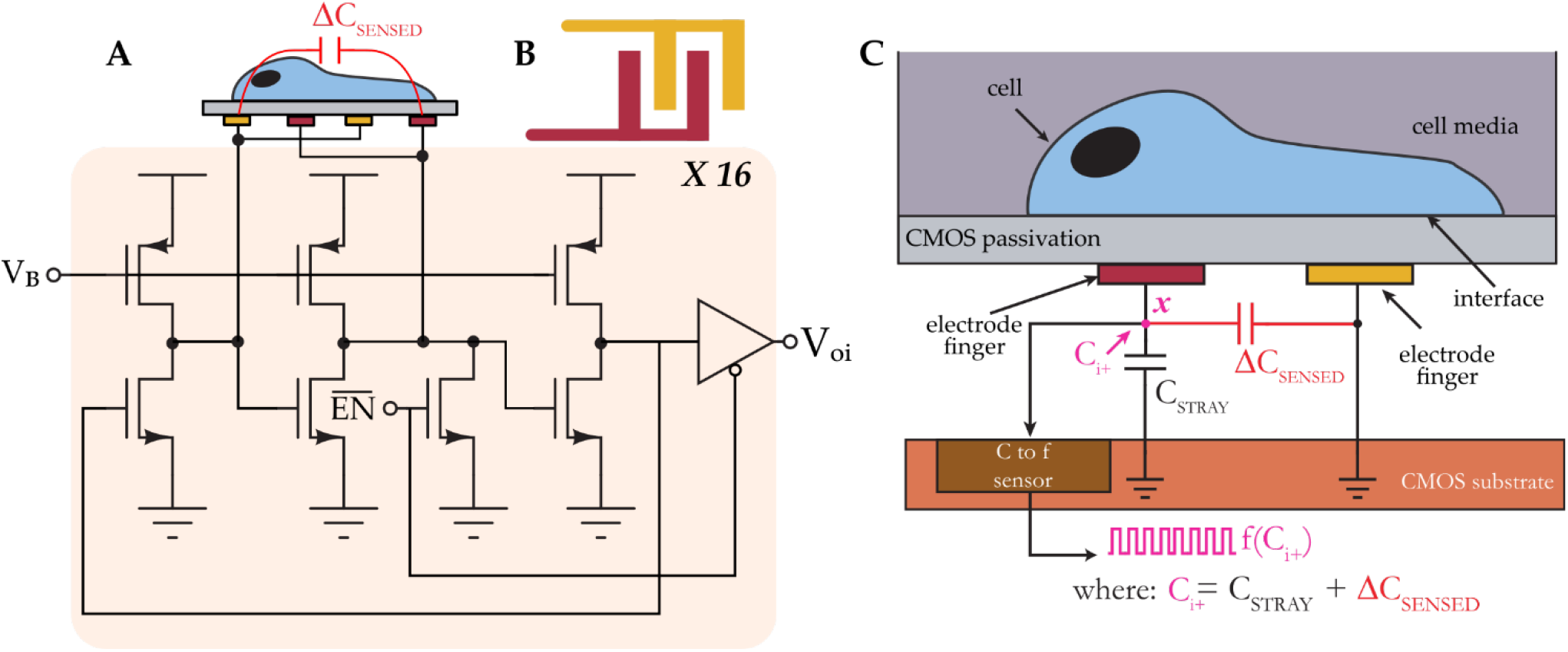
(A) System architecture of a CMOS capacitance sensor. The sensor consists of 16 individually-addressable capacitance-to-frequency pixels, and each pixel includes a ring oscillator circuit whose base frequency can be tuned via a bias current established by the voltage V_B_. An interdigitated passivated electrode structure is connected to the intermediate stage of the oscillator. Each pixel can be turned off independently using an enable signal (EN_BAR), and each pixel outputs a digital signal V_oi_ when turned on. (B) Top view of an interdigitated electrode. (C) Cross-sectional view showing the sensor’s transduction mechanism. The electrode fingers establish a coupling field at the interface, and perturbations in the field due to cell adsorption, movement, or life cycle events modulate the electrode’s effective capacitance. This change is mapped to the frequency of a test signal using a capacitance-to-frequency sensor.

The input capacitance to the electrode is labeled ΔC_SENSED_, and it consists of two capacitances. The first is the electrode’s intrinsic capacitance, which is variable, and the second is a fixed parasitic capacitance that exists between the electrode and a reference node of the circuit. The effective capacitance C_i+_ at node ***x***, which consists of the parallel combination of the two aforementioned capacitances, is mapped to a frequency f(C_i+_). In a cell assay using this sensor, for example, during the sedimentation phase when a cell gets close to the interdigitated electrode, ΔC_SENSED_ changes from its baseline value as a result of perturbations in the coupling field established by the electrode fingers; these perturbations change the electrode’s intrinsic capacitance. The frequency of the test signal is continually monitored, and it is used to calculate a relative change in capacitance, and this change in capacitance is subsequently correlated with cell activity.

### 2.2 System Integration and Experimental Setup

The microsystem included an integrative package capable of maintaining the chip’s electrical integrity while allowing liquid samples to be applied onto its sensing surface. The packaging procedure employed was described in previous publications (Dandin *et al*., 2009; Datta-Chaudhuri, Abshire and Smela, 2014; Gilpin *et al*., 2022). Briefly, it included attaching the ASIC die directly onto a PCB daughter board, which included a redistribution pad frame. Once attached to the PCB, the die’s I/O pads were wire-bonded directly to the leads of the redistribution pad frame. The wire-bonds were encapsulated with an impermeable epoxy. A subsequent encapsulation step was employed, extending the encapsulant from the edge of the chip to the edge of the redistribution platform in order to make a platform onto which additional structures could be built. A cell culture-compliant dish was then glued to the resulting epoxy platform. Within the cell culture dish, a second chamber was made using a 1 cm spectroscopy cuvette. Two holes were machined at the base of the cuvette in order to allow fluid exchange between the cuvette and the outer dish. The cells were plated inside the cuvette, and cell media was used to fill the dish with enough fluid to cause the cuvette to be filled to the brim, at which point a glass cover slip was placed on top of the cuvette. This two-chamber arrangement was used in order to minimize fluid evaporation over the sensing area as well as to maintain a constant focus for imaging (B. Senevirathna, Lu, Smela, *et al*., 2019).

All experiments were conducted in a cell culture incubator at 37° C and under 5% CO_2_. A bright field upright optical microscope was placed inside the incubator to monitor the cells on top of the chip. Microscopy images were obtained every 5 minutes over the duration of the experiment (typically 24 to 48 hours) using a C-mount digital camera attached to the microscope. The data acquired from the chip and the images were automatically uploaded to a decentralized cloud environment where image processing and data analytics were performed using custom algorithms. Figure 2 illustrates the various components of the system.

**Figure 2.**
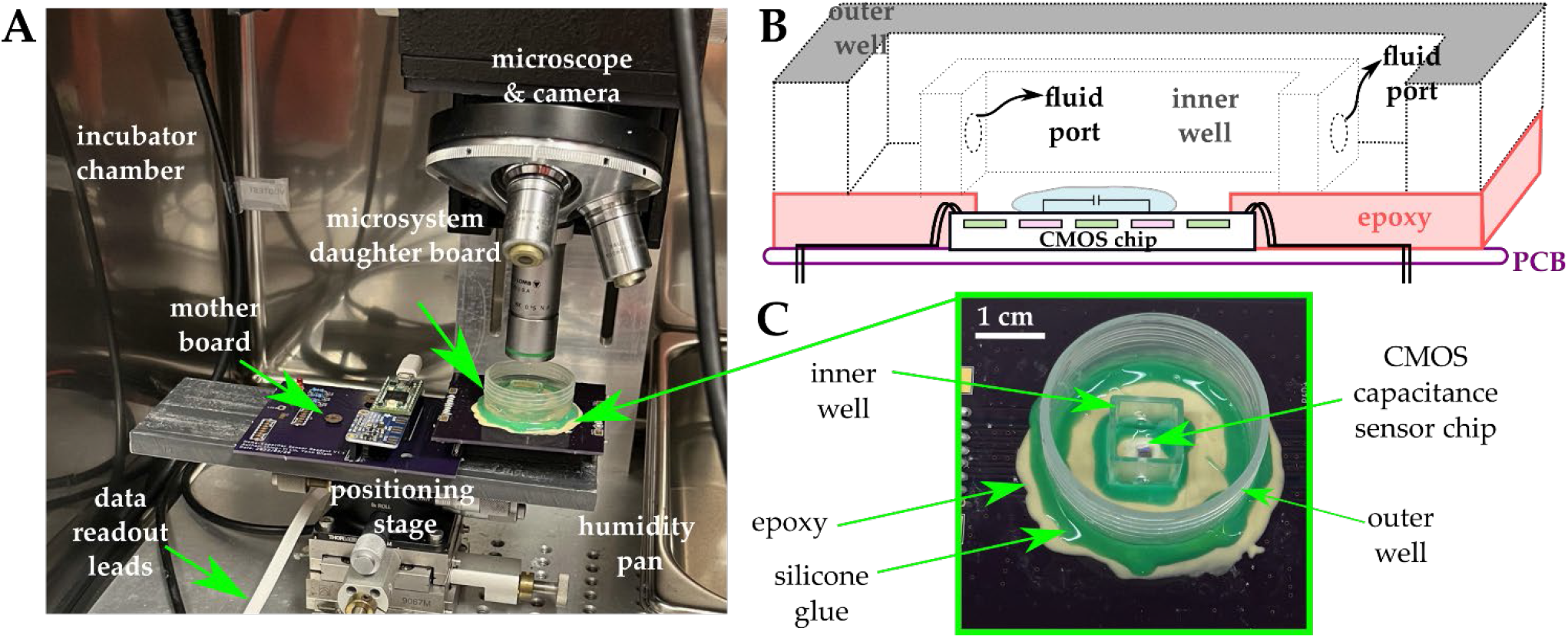
(A) The experimental setup includes an incubator in which a microscope and a camera are mounted. The microsystem is mounted on a daughter board and subsequently connected to a mother board that includes a microcontroller and additional ancillary circuits for data acquisition and control. Positioning is achieved using a micro-positioning stage. Data from the chip and from the camera are read out via leads that are implemented using flat USB cables to allow the incubator door to close and maintain a sealed environment. Humidity pans are also used for additional environmental control. (B) Cross-sectional view of the integrative package used in the study. A two-well system is used in order to maintain a constant focus and minimize evaporation. (C) Photograph of the microsystem.

### 2.3 Cell Culture

Murine Balb/c Raw 264.7 macrophages (ATCC) were cultured in a T75 flask with culture media containing DMEM (ThermoFisher Scientific) without phenol red and with HEPES with 10% FBS (VWR). Culture media was removed from the T75 flask, and 7 mL of sterile PBS was added to the culture flask and incubated for 5 minutes to wash the cells. The PBS was removed, and 7 mL of media was added to the flask. A cell scraper (VWR) was used to resuspend the cells in the media. A volume of 5 mL containing the cells was added to a 15 mL centrifuge tube and centrifuged for 5 minutes. The supernatant was removed, 2 mL of fresh media was added to the centrifuge tube, and the cells were resuspended and added to a new T75 flask with 8 mL of fresh media. From the estimated cell density, the necessary volume was pipetted so that the starting number of cells seeded in the microsystem was ~2 million. We note that upon seeding the cells in the microsystem’s analysis well, only a small fraction of the starting cell population fell in the vicinity of the electrode array.

### 2.4 Data Collection and Analysis

The data yielded by the experiments consisted of a time series dataset originating from measurements from the 16 sensors comprised on the chip and of an imaging dataset originating from micrographs acquired by the camera attached to the microscope. The time series data consisted of capacitance change measurements (ΔC) from all the electrodes, performed every 29 seconds. As noted previously, the imaging dataset was generated by taking an image of the cell culture every 5 minutes. Both datasets were time-stamped automatically by the control program, and experiments were typically conducted for 48 hours. Exemplary data from each data set are shown and discussed in detail in the following subsections.

#### 2.4.1 Imaging Dataset

Figure 3 shows a subset of images from the imaging dataset, with a white box enclosing the region of interest (ROI), *i.e*., the sensing area spanned by the 4 × 4 electrode array. The images were post-processed using image processing packages available in python (OpenCV, matplotlib, and numpy). Post-processing included contrast enhancement, sharpening, and applying a false color scheme. These post-processing steps were conducted to enhance image quality in order to better visualize the cells adhered to microchip’s surface and further to facilitate the automatic detection and counting of cells using a custom-designed Python algorithm for cell identification and pattern classification. This algorithm is further described in detail below.

**Figure 3.**
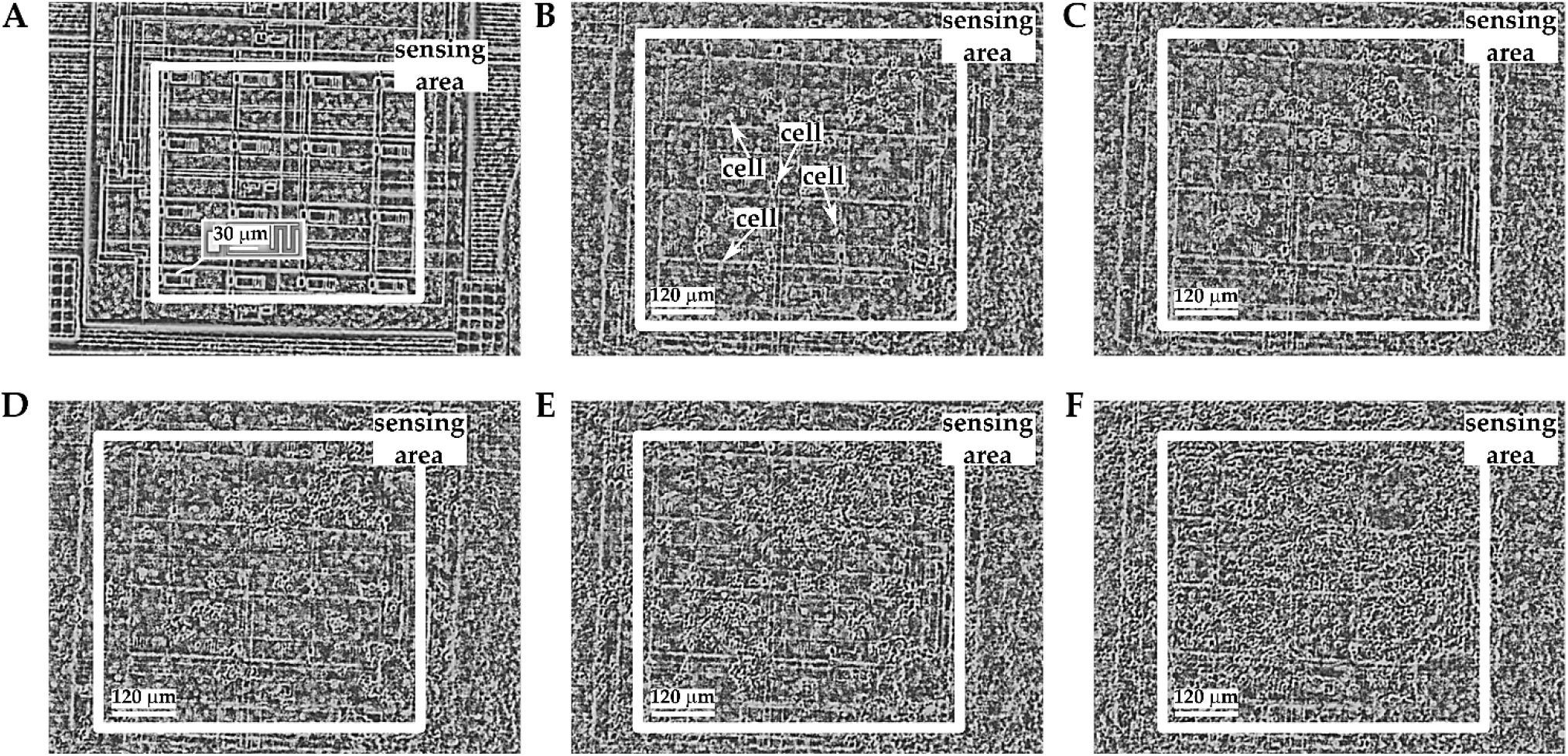
(A) Photomicrograph of a microchip without cells plated thereon. The sensing area is shown by the white box, which encloses the 4 × 4 electrode array. The inset shows a photomicrograph of the interdigitated electrode used in the pixel array. (B)-(F) Photomicrographs of the microchip when macrophages are plated thereon, at time *t* = 0, *t* = 8 hours, *t* = 15 hours, *t* = 28 hours, and *t* = 44 hours. Cells are identified with the white arrows in panel B. Note: the photomicrograph in panel (A) is at a different scale than the photomicrographs in panels (B)-(F).

Cell counting was effected using a custom template matching algorithm developed in Python. Briefly, the algorithm first included pre-processing the images of the dataset, as described above. Second, using a reference image with cells present in the ROI, a template was extracted by overlapping each cell in the image with its center aligned to the others and taking an average intensity. The template served as a correlation filter which the algorithm attempts to retrieve in subsequent images in the dataset. When a high correlation between a feature in an image and a template was obtained, the algorithm flagged the location as likely having a cell, and it marked the image with a graphical indicator (*e.g*., a red circle, as shown in Figure 4) at that location (Kim and de Araújo, 2007; Bolme, Draper and Beveridge, 2009). To count the cells in the image, the algorithm reported the number of indicators registered upon parsing the ROI on a pixel-by-pixel basis. A result of the cell counting algorithm is shown in Figure 4 for four representative images.

**Figure 4.**
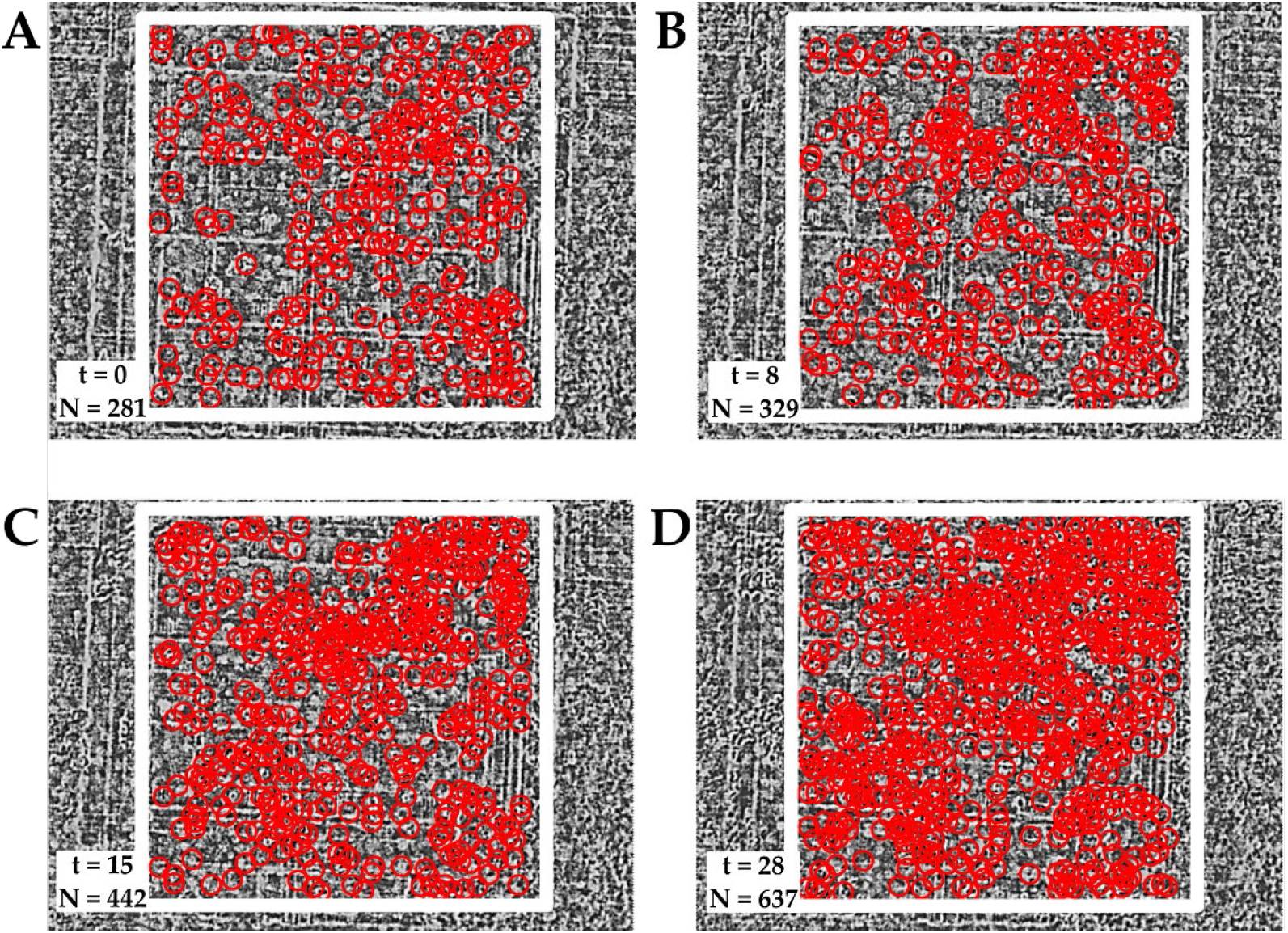
Results of the cell counting algorithm showing the time *t* at which a photomicrograph was taken and the number cells *N* that were estimated to be in the area. The red circles indicate areas that are likely occupied by a cell. Panels A, B, and C show the data (*t, N*) for four different time points in a macrophage culture experiment. The white box delineates the ROI.

#### 2.4.2 Time Series Dataset

The time series data set included measurements from the capacitance sensors comprised in the array. A measurement was taken every 29 seconds, yielding 16 data points (one for each electrode), and for the duration of the experiment. The change in capacitance was calculated by subtracting an initial measurement of capacitance prior to the start of the experiment from each of the instantaneous measurements obtained during the experiment. Figure 5A shows the resulting measurement traces across the array for the macrophage growth experiment depicted in Figure 3.

**Figure 5.**
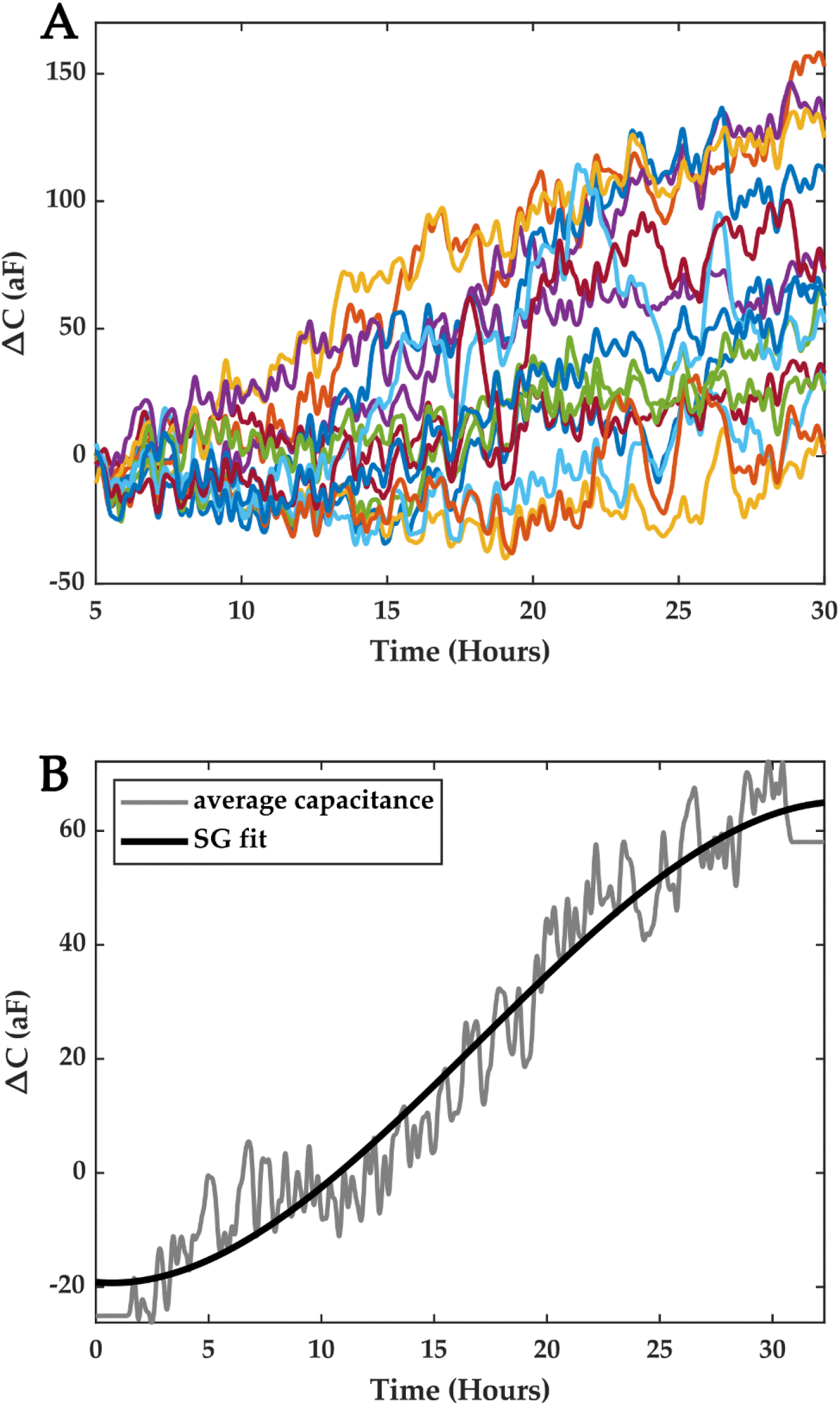
(A) Time series data obtained from a macrophage growth experiment. The traces corresponds to the measurements from the sensor array. (B) Average capacitance data (gray) were obtained by averaging the measurements from all the pixels. The solid black trace is a trend-preserving fit that is obtained using a Savitsky-Golay filter.

Contrary to previous works on integrated capacitance data for cell measurements where cell coverage of the electrode was correlated with the measured capacitance, in this study, we consider the average capacitance obtained from all the electrodes in the array and seek to correlate it with cell *growth*. We argue that this average capacitance can be considered as an overall indicator of cell growth in the ROI, where each electrode serves to sample the ROI spatially, and we seek to establish a temporal model that links cell numbers and the average capacitance.

Furthermore, to obtain a capacitance curve that characterizes capacitance growth in the ROI for a given period of time, we utilize the trend of the data rather than instantaneous average capacitance data. This means local temporal variations in capacitance due to Brownian motion, cell movements in the vicinity of the electrodes, and detector noise are averaged out. To obtain this trend, we fitted the average data with a Savitsky-Golay (SG) filter of the third order, with a frame length equal to *n-1*, where *n* is even and is the number of points in the average capacitance time series (Savitzky and Golay, 1964). Figure 5B shows the results of the fit, which were obtained using Matlab. This fit captures the trend of the capacitance registered in the ROI over the course of an experiment for a period of ~ 30 hours.

To compute a single capacitance growth metric, one may simply perform a piece-wise linear approximation of the SG fit and estimate therefrom a capacitance growth factor *S* (in ΔC/hour) for each segment, *S* being the slope of the segment. An average capacitance growth parameter *S_avg_* may be computed by averaging all the slopes. The piece-wise linear approximation may be performed by dividing the SG trace in 1-hour overlapping segments, with a half-hour overlap between segments. Subsequently, a linear regression may be conducted on each segment to estimate the average capacitance growth factors *S*, from which Savg may be computed. Alternatively, a histogram of all the slopes calculated from SG fits of each pixel trace may be constructed, and the mean value of the resulting distribution can be used as the average capacitance growth parameter *S_avg_*. The standard deviation of the distribution provides an appreciation for the deviation away from the mean capacitance growth factor registered during the experiment.

The latter method was used, and its results are shown in Figure 6. The capacitance growth factors obtained for the entire dataset were binned to form a histogram that highlighted the diversity of capacitance growth factors across the ROI, and the mean of this distribution provided an average capacitance growth factor estimate over the ROI, denoted *S_avg_*. For the featured dataset, the average capacitance growth factor *S_avg_* was 3.38 aF/hr, with a standard deviation of 16.57 aF/hr, and the distribution was assumed to be Gaussian.

**Figure 6.**
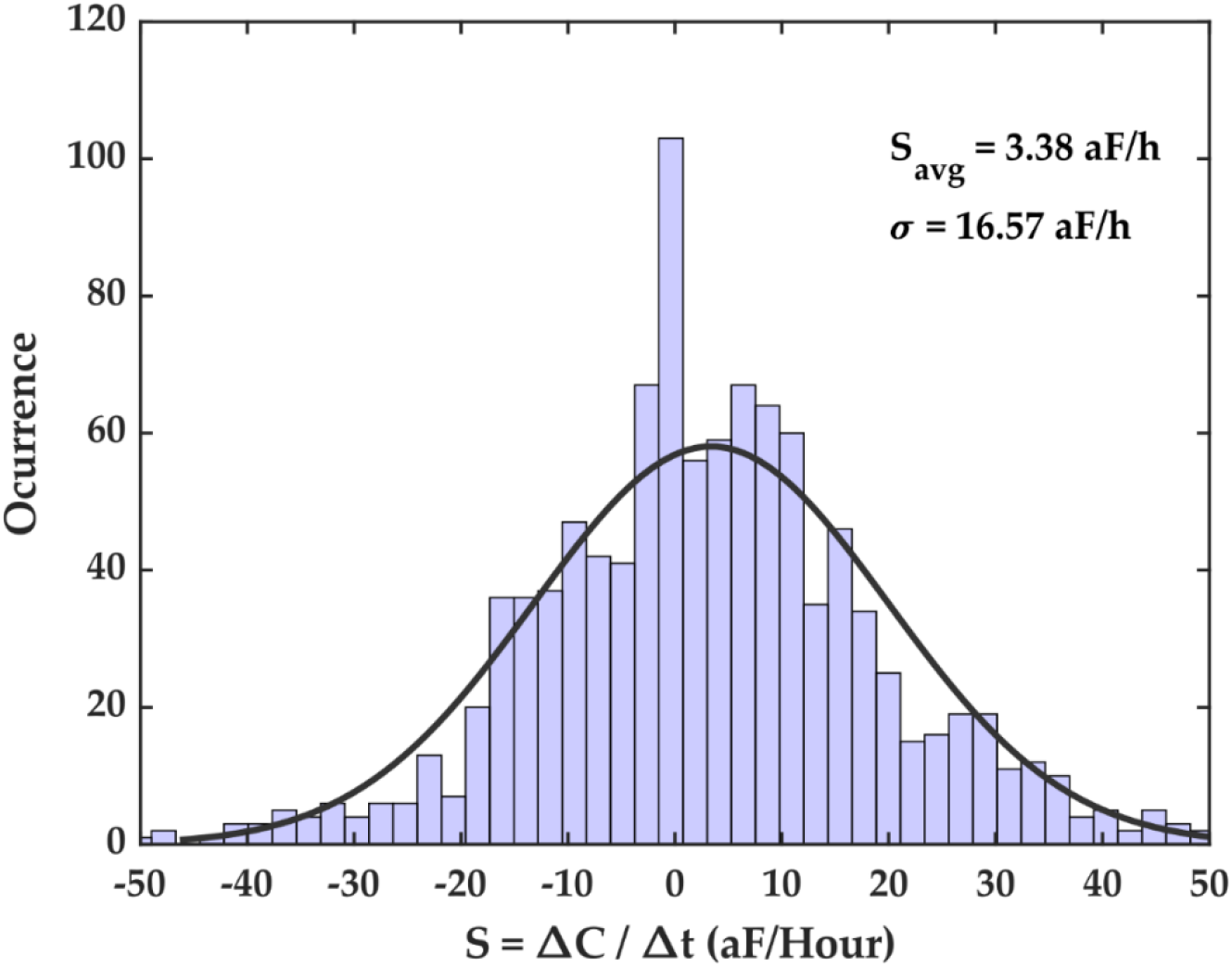
Capacitance growth factor (S) histogram for the time series capacitance dataset. This histogram shows the extent of capacitance growth factors over the entire electrode array as well as the number of occurrences of measured capacitance growth factors in the dataset. The distribution’s mean is *S_avg_* = 3.38 aF/hour, with a standard deviation of *σ* = 16.57 aF/hour. The parameter *S_avg_* is the average capacitance growth registered during the experiment.

## 3 Results

We conducted several growth experiments and analyzed their results using the methods discussed above. We feature herein three representative experiments where macrophages were cultured on the chip and left to proliferate for 48 hours. The first 30 hours of each experiment were considered for studying macrophage growth dynamics and for correlating measured capacitance results with cell counts estimated via imaging. This time frame was chosen empirically based on observations that the ROI would saturate with cells beyond 30 hours, thus resulting in a reduced cell growth rate in the ROI. Figure 7 illustrates the average SG fits for each of the three experiments starting from t = 0 to t = 30 hours.

**Figure 7.**
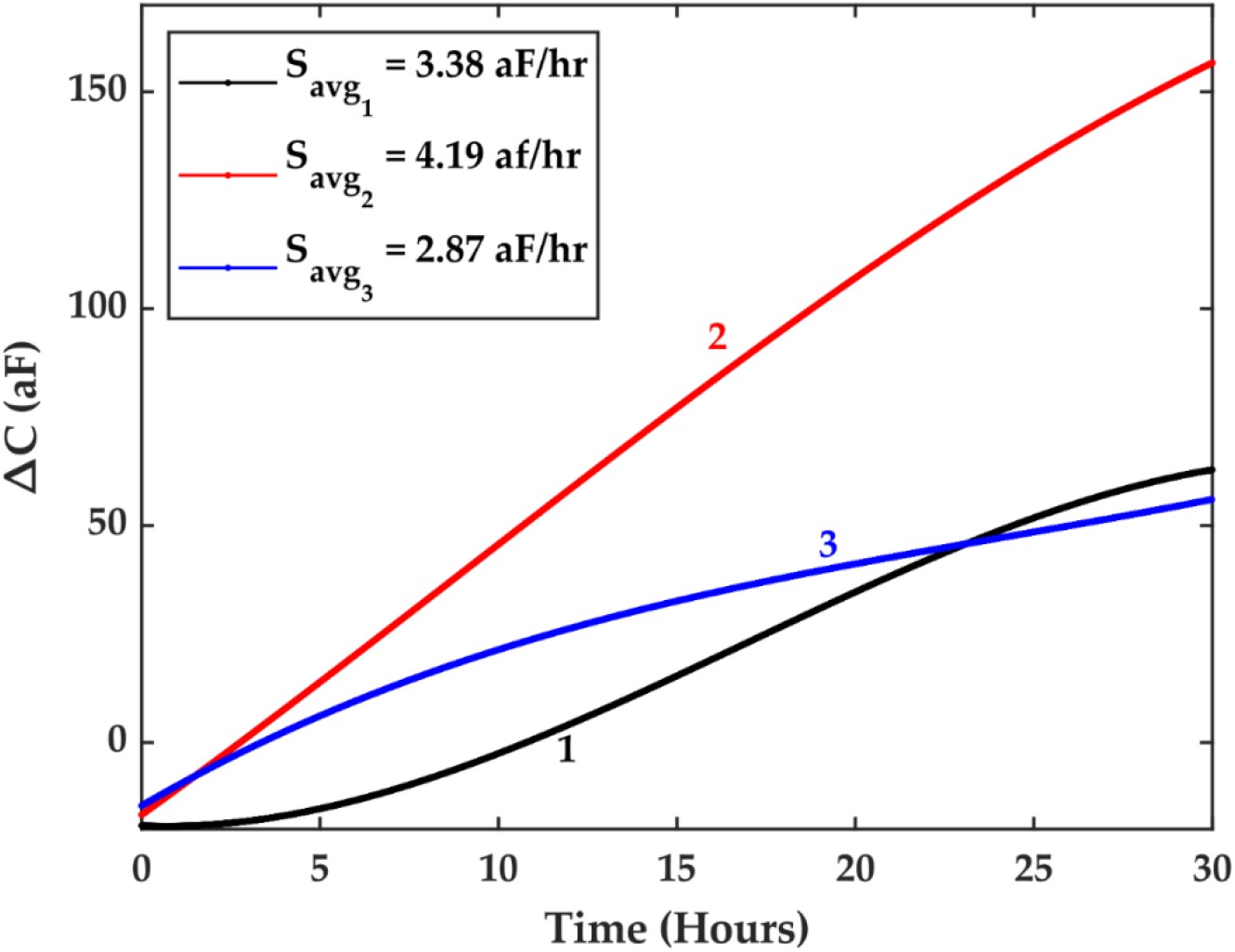
SG trends for three different experiments. The growth phase is assumed to be within the first 30 hours. Different average capacitance growth factors *S_avg_* were registered across the three experiments.

Cell counts were estimated for each experiment using our custom Python code. To do so, a subset of images from the imaging dataset was chosen. The time stamps for each image were extracted, and the number of cells in the ROI was estimated by the algorithms. Furthermore, the values of the average capacitance data at the same time stamps were extracted from the time series dataset. Our results show that the natural logarithm of the cell counts was highly correlated with the average capacitance and that the correlation was linear (Figure 8).

**Figure 8.**
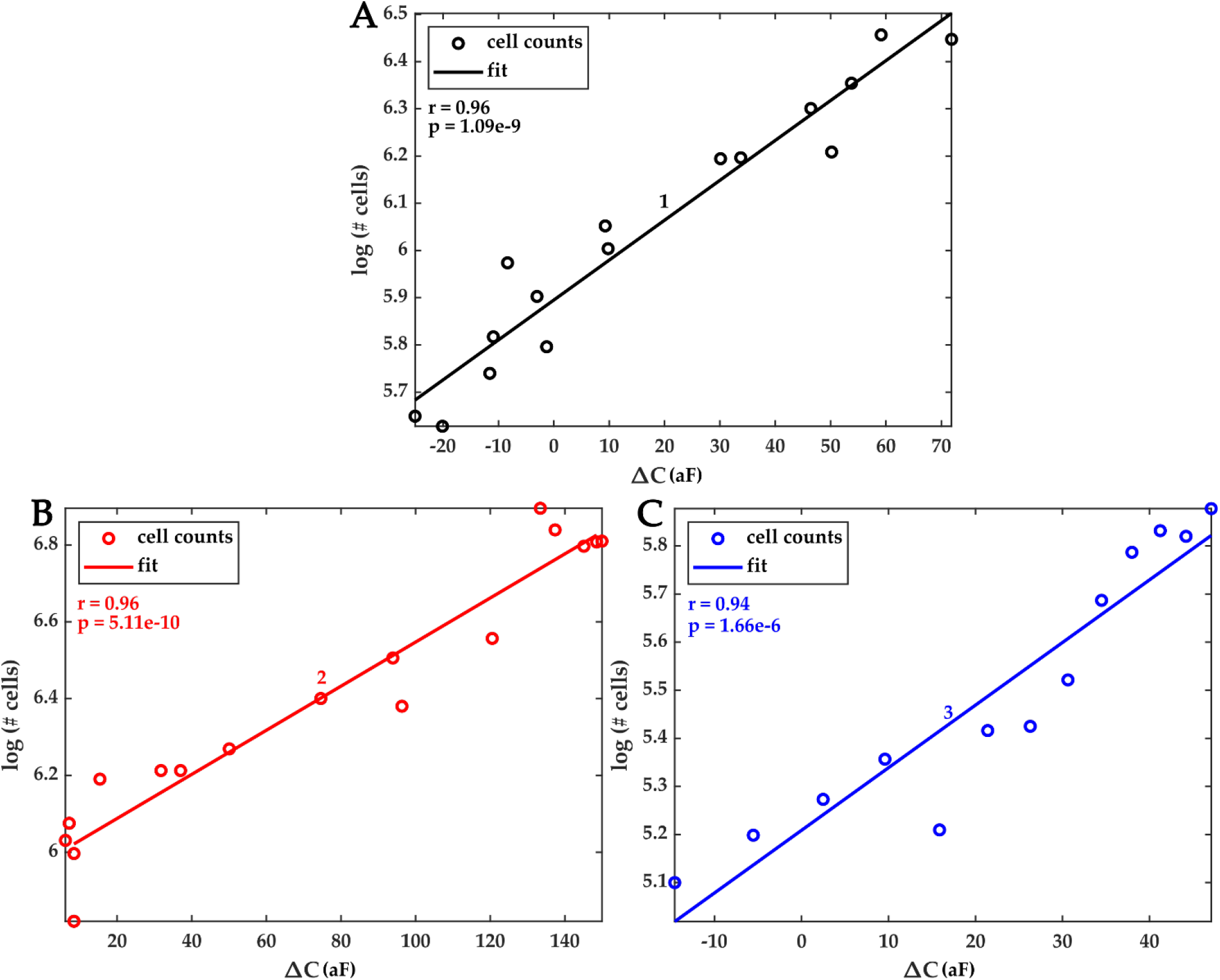
Correlation plots of estimated cell counts (in natural logarithm scale) and the measured average capacitance. (A)-(C) Pearson’s correlation coefficients *r* and its corresponding *p*-value for the three featured experiments (1, 2, and 3) were (r = 0.96, p-value < 10^-8^), (r = 0.96, p-value < 10^-9^), and (r = 0.94, p-value < 10^-5^), respectively.

This finding, namely that the logarithm of cell counts and average capacitance follow a linear relationship, lays the foundation for a time-dependent model that can be used to predict future cell numbers or to calculate the instantaneous number of cells in the ROI, after sufficient capacitance measurements and images have been collected.

To construct a temporal model that relates cell numbers and capacitance, we first used a linear regression to fit a line between the logarithm of the measured number of cells and the measured average capacitance data for the ROI, and we write Equation 1, which captures this log-linear relationship. Here, *N_m_* is the number of cells, *α* is the slope of the line, *ΔC_m_* is the average capacitance of the ROI, and *K* is the intercept of the line.

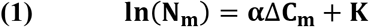

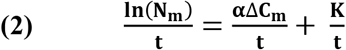

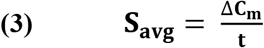

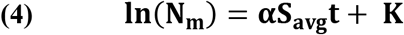

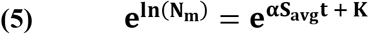

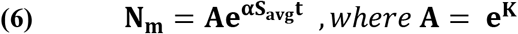

Equation 1 is normalized by the variable *t*, yielding Equation 2, which introduces the time dependence in the model. Here, the implicit assumption that is made is that the log-linear relationship between the number of cells and the measured average capacitance is itself time-invariant. Specifically, we assume that neither α nor *K* has a time dependence. We will discuss in the following section situations under which this assumption fails and in which, consequently, the proposed model cannot accurately capture the temporal evolution of the cell population within the ROI.

In Equation 3, we assume that the term *ΔC_m_/t* is the average capacitance growth factor *S_avg_*. As previously noted, the *S_avg_* factor is a measure of the average rate of change of the capacitance for a normally distributed set of average rates across the ROI. Thus, our assumption here is justified since that for any given capacitance growth rate *ΔC_m_/t*, there is a roughly 70% probability that this rate is within one standard deviation of the mean of the distribution. This, of course, is because the distribution is assumed to be Gaussian, as shown in Figure 6. Lastly, there is another caveat. For distributions with large standard deviations, Equation 2 is less likely to hold. However, we posit that datasets with large standard deviations would fall in the category for which the model would not hold anyway, as we shall discuss later.

Equations 4 and 5 show the intermediate steps that lead to Equation 6, which broadly states that the number of cells in the ROI follows an exponential dependence time, and where the argument of the exponential includes experimentally determined constants *α*, *A*, and *S_avg_*. Predictably, simply fitting Equation 6 to the measured cell counts as a function of time does not work. This is readily observable when one considers that the constant *A* depends on the y-intercept of the linear fit. As such, at t = 0, Equation 6 is likely to predict the cell count erroneously. In other words, the intercept may differ enough from the actual value and bias the exponential away from the measured data.

Thus, we posit that Equation 6 is a generalized temporal model that links measured capacitance growth factor and measured cell numbers. Consequently, Equation 6 must be updated further in order to obtain an experiment-specific model that depends on independent variables that are particular to the experiment. These independent variables consist of time, which is already captured in Equation 6, and two other independent variables which are the cell numbers at two specific times. The latter two variables form boundary conditions for Equation 6. The first boundary condition is *N_0_, i.e*., the number of cells at time *t_0_*, and the second boundary condition is *N_f_*, the number of cells at time *t_n_*, where *n* is strictly positive. The particular model is formulated in Equation 7. The constants *Γ* and *β* can be determined using the boundary conditions shown in Equations 8 and 9, provided *N_0_* and *N_f_* are known. The parameters *α* and *S_avg_* are determined experimentally using the imaging and time series datasets for the time interval [*t_0_, t_n_*].

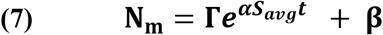

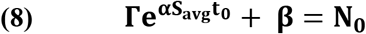

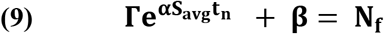

We tested the particular temporal model against the cell count measurements for each of the three experiments. In each case, good agreement was found between the estimated cell counts, as can be seen in Figures 9A–9C. The goodness of the model was evaluated using an adjusted-R^2^ coefficient of determination, where the number of independent variables was *k* = 3.

**Figure 9.**
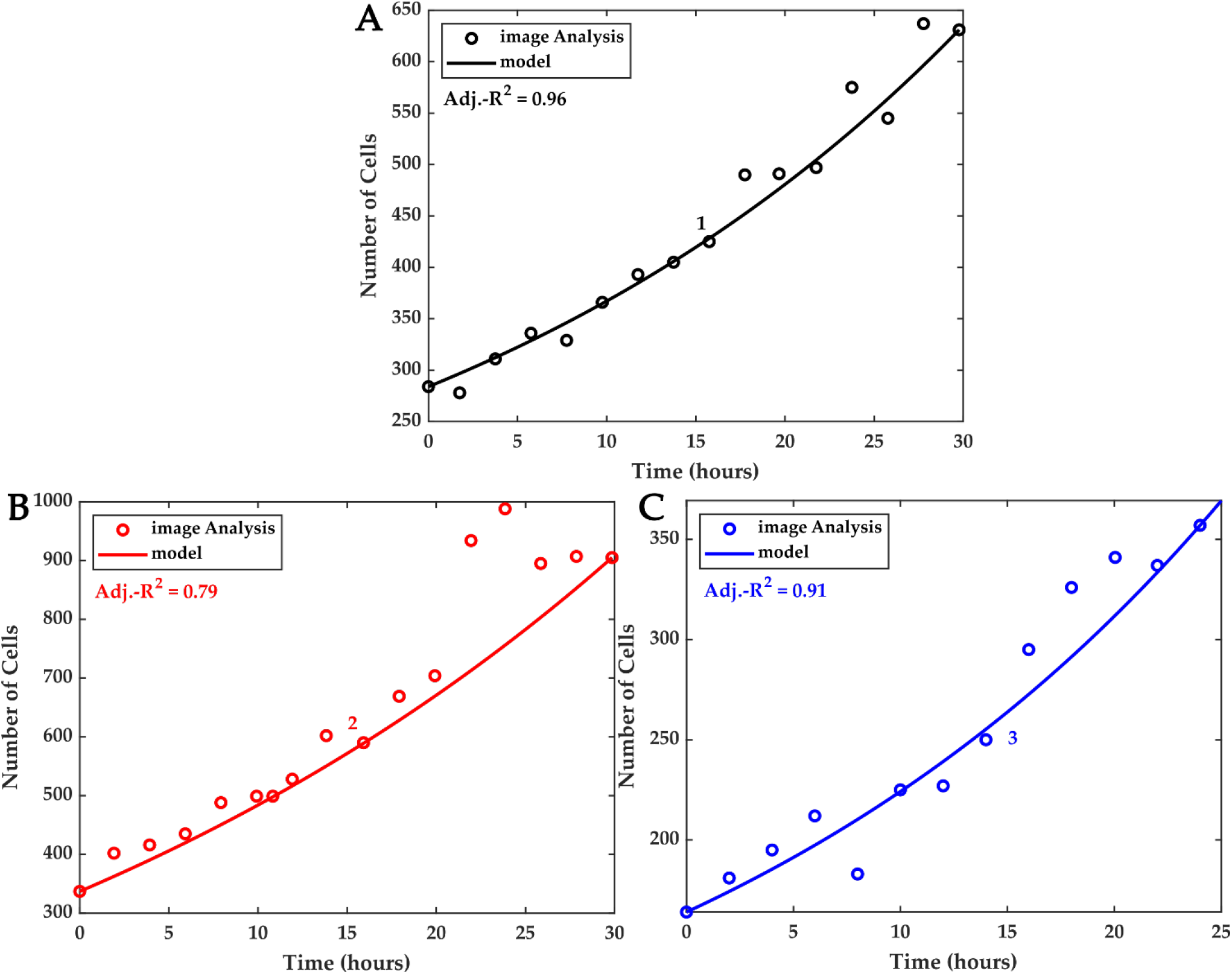
Temporal model linking cell numbers and capacitance. The model utilizes the average measured capacitance growth *S_avg_* to account for the cell numbers at time *t* ∈ [*t_0_* = 0, *t_f_* = 30] for Experiments 1 and 2 (shown in panels (A) and (B)), and at time *t* ∈ [*t_0_* = 0, *t_f_* = 25] for Experiment 3 shown in panel (C).

Table 1 shows the measured results for the experiments. We report the *S_avg_* parameters computed for each of the three experiments and the Pearson correlation coefficient and its associated p-value, which assess the correlation between measured capacitance and measured cell numbers. Further, we report the R^2^ and the adjusted R^2^ values of the proposed model. Table 1 also reports the doubling time calculated from the model.

**Table 1.** Summary of experimental results and statistics for the three experiments. The last column of the table shows the estimated doubling time *t*_d_ based on the proposed model.

| *Experiment ID* | $S_{avg}$<br>(aF/hr) | $\sigma$<br>(aF/hr) | $r_{c,n}$ | *p-value* | $R^2$ | *adj.R<sup>2</sup>* | $t_d$ |
| --- | --- | --- | --- | --- | --- | --- | --- |
| *1* | 3.38 | 16.57 | 0.96 | 1.09E-9 | 0.97 | 0.96 | 26 |
| *2* | 4.19 | 20.43 | 0.96 | 5.11E-10 | 0.83 | 0.79 | 20 |
| *3* | 2.87 | 25.15 | 0.94 | 1.66E-6 | 0.93 | 0.91 | 21 |

For comparison, the reported average doubling time of the macrophages used in our study varies from 12 hours to 15 hours, depending on the experimental conditions used (Bancos and Tyner, 2014; Jarrar, Çetin Altindal and Gümüşderelioğlu, 2020). In contrast, our results showed an average doubling time of 22.3 hours, and this figure was consistent with the growth rate measured for the same cells in our lab using standard cell counting procedures.

## 4 Discussion and Conclusions

This study demonstrated a system configured for tracking macrophage growth *in situ* during their culture on an electronic chip. While cell culture on electronic chips has been shown before, particularly on capacitance sensing chips, several novel contributions stem from our work. We demonstrate that a set of electrodes that are sparsely disposed within a region of interest can serve to monitor cell growth kinetics temporally over the entire region. Specifically, we show that an average capacitance growth factor measured by all the electrodes in the area is a linkage factor for a model that relates cell numbers to measured capacitance. This model (shown in Equations 7–9) accurately captures cell growth, provided that the number of cells is known at two distinct times during the growth phase.

One of the model’s key determinants is the degree of correlation that exists between the measured cell counts and the average capacitance when estimating the parameter *α* in Equation 7. Recall that this parameter is the rate of change of the number of cells as a function measured capacitance, and we assumed that it was constant and time-invariant, and thus that capacitance and cell numbers were linearly correlated. These conditions must be met in order to ensure that the temporal model accurately describes cell growth in the ROI.

However, there are several conditions under which the degree of correlation between average capacitance and cell numbers may be reduced, which would reduce the accuracy of the model shown in Equations 7–9. For instance, when the ROI experiences non-negligible cell migration, whether in or out of the ROI, there may be a non-linear correlation between measured capacitance and measured cell numbers. This scenario would likely invalidate the assumption that *α* is constant time-invariant, and Equations 7–9 would thus not be adequate to describe the cell dynamics within the ROI. Such a scenario may cause a larger spread around a mean capacitance growth factor, since electrodes that are closer to the migration sites in the ROI may experience a much greater or much lower average growth factor, depending on the direction of the migration.

In yet other situations, the correlation between the two measured variables may be reduced. For example, the degree of sparsity of the electrodes inside the ROI may significantly reduce the correlation between the observed capacitance and the number of cells inside the ROI. This is because the farther apart the electrodes are, the more localized their individual responses are. As such, measured capacitance would only correlate with small groups of cells that are in the vicinity of the electrodes. Conversely, in ROIs that are densely populated with electrodes, it is more likely to obtain high correlations, and thus, this should be a design goal in future capacitance-sensing lab-on-CMOS devices. However, increasing electrode density has practical implications for performance: the closer electrodes are placed, the more a single electrode serves as a parasitic load to its nearest neighbors. This would reduce electrode sensitivity. As such, there is a density-to-sensitivity trade-off that must be mitigated at the circuit design phase.

Lastly, the techniques shown herein may be used as part of a digital signal processing (DSP) and sensing framework for lab-on-CMOS capacitance sensors. For instance, the proposed model may be used periodically and selectively, *i.e*., on chosen time segments during data acquisition, to predict short-term cell populations swings. Such information could then be used to issue control signals to release drugs or other external stimuli in the cell media in order to determine their effects on cell culture kinetics. Generally, the predictive analyses enabled by this framework will permit the monitoring of cell cultures *in situ* with much greater temporal resolution and parallelism than what is currently possible with the state-of-the-art.

## 6 Author Contributions

KS conducted the macrophage growth experiments and analyzed the data. He further developed the model for correlating measured capacitance and cell growth, and he wrote the original manuscript. CYL developed analytical methods for the time series data and for image processing, and he designed the information infrastructure for cataloging measured data into the Integrated Circuits and Bioengineering laboratory’s dedicated Oracle™ cloud environment. Further, CYL designed the mother board. YG developed the integrative packaging method and designed the daughter board. He also developed the code for data acquisition and control, and he built the experimental fixture used inside the incubator. YG further trained and supervised KS in his initial efforts using the lab-on-CMOS platform. EW supervised the macrophage growth experiments. MD supervised the development of the experimental setup, the microsystem design, and he further supervised technology integration efforts as well as the experiments, data collection, and data analysis. EW and MD designed the study, secured funding, and provided resources and guidance to KS, CYL, and YG throughout the study. KS, EW, CYL, and MD contributed text and figures to the manuscript and edited it.

## 8 Data Availability

The datasets generated for this study are available upon request to the corresponding authors.

## References

Abdelhamid, H. et al. (2022) ‘A capacitive sensor for differentiation between virus-infected and uninfected cells’, Sensing and Bio-Sensing Research, 36, p. 100497. doi:10.1016/j.sbsr.2022.100497.

Bancos, S. and Tyner, K.M. (2014) ‘Evaluating the effect of assay preparation on the uptake of gold nanoparticles by RAW264.7 cells’, Journal of Nanobiotechnology, 12(1), p. 45. doi:10.1186/s12951-014-0045-5.

Bolme, D.S., Draper, B.A. and Beveridge, J.R. (2009) ‘Average of Synthetic Exact Filters’, in 2009 IEEE Conference on Computer Vision and Pattern Recognition. IEEE, pp. 2105–2112. doi:10.1109/CVPR.2009.5206701.

Bunnfors, K. et al. (2020) ‘Nanoparticle activated neutrophils-on-a-chip: A label-free capacitive sensor to monitor cells at work’, Sensors and Actuators B: Chemical, 313, p. 128020. doi:10.1016/j.snb.2020.128020.

Christen, J.B. and Andreou, A.G. (2007) ‘Design, fabrication, and testing of a hybrid CMOS/PDMS microsystem for cell culture and incubation’, IEEE Trans. Biomed. Circuits Syst., 1(1), pp. 3–18. doi:10.1109/TBCAS.2007.893189.

Dandin, M. et al. (2009) ‘Post-CMOS packaging methods for integrated biosensors’, in Proceedings of IEEE Sensors. doi:10.1109/ICSENS.2009.5398540.

Datta-Chaudhuri, T., Abshire, P. and Smela, E. (2014) ‘Packaging commercial CMOS chips for lab on a chip integration’, Lab on a Chip, 14(10), p. 1753. doi:10.1039/c4lc00135d.

Derlindati, E. et al. (2015) ‘Transcriptomic Analysis of Human Polarized Macrophages: More than One Role of Alternative Activation?’, PLOS ONE, 10(3), p. e0119751. doi:10.1371/journal.pone.0119751.

Ferlito, U. et al. (2020) ‘Sub-Femto-Farad Resolution Electronic Interfaces for Integrated Capacitive Sensors: A Review’, IEEE Access, 8, pp. 153969–153980. doi:10.1109/ACCESS.2020.3018130.

Forouhi, S., Dehghani, R. and Ghafar-Zadeh, E. (2018) ‘Toward high throughput Core-CBCM CMOS capacitive sensors for life science applications: A novel current-mode for high dynamic range circuitry’, Sensors, 18(10). doi:10.3390/s18103370.

Forouhi, S., Dehghani, R. and Ghafar-Zadeh, E. (2019) ‘CMOS based capacitive sensors for life science applications: A review’, Sensors and Actuators, A: Physical, 297, p. 111531. doi:10.1016/j.sna.2019.111531.

Ghallab, Y.H. and Ismail, Y. (2014) ‘CMOS Based Lab-on-a-Chip: Applications, Challenges and Future Trends’, IEEE Circuits Syst. Mag., 14(2), pp. 27–47. doi:10.1109/MCAS.2014.2314264.

Gilpin, Y. et al. (2022) ‘Tracking the Effects of Tumor Treating Fields on Human Breast Cancer Cells in vitro Using a Capacitance Sensing Lab-on-CMOS Microsystem’, in 2022 29th IEEE International Conference on Electronics, Circuits and Systems (ICECS). IEEE, pp. 1–4. doi:10.1109/ICECS202256217.2022.9971093.

Hedayatipour, A., Aslanzadeh, S. and McFarlane, N. (2019) ‘CMOS based whole cell impedance sensing: Challenges and future outlook’, Biosens. Bioelectron., 143(June), p. 111600. doi:10.1016/j.bios.2019.111600.

He, L. et al. (2021) ‘Global characterization of macrophage polarization mechanisms and identification of M2-type polarization inhibitors’, Cell Reports, 37(5), p. 109955. doi:10.1016/j.celrep.2021.109955.

Hu, K., Arcadia, C.E. and Rosenstein, J.K. (2021) ‘A large-scale multimodal CMOS biosensor array with 131,072 pixels and code-division multiplexed readout’, IEEE Solid-State Circuits Lett., 4, pp. 48–51. doi:10.1109/LSSC.2021.3056515.

Jarrar, H., Çetin Altindal, D. and Gümüşderelioğlu, M. (2020) ‘The inhibitory effect of melatonin on osteoclastogenesis of RAW 264.7 cells in low concentrations of RANKL and MCSF’, Turkish Journal of Biology, 44(6), pp. 427–436. doi:10.3906/biy-2007-85.

Kim, H.Y. and de Araújo, S.A. (2007) ‘Grayscale Template-Matching Invariant to Rotation, Scale, Translation, Brightness and Contrast’, in, pp. 100–113. doi:10.1007/978-3-540-77129-6_13.

Konakovsky, V. et al. (2015) ‘Universal Capacitance Model for Real-Time Biomass in Cell Culture’, Sensors, 15(9), pp. 22128–22150. doi:10.3390/s150922128.

Li, L., Yin, H. and Mason, A.J. (2018) ‘Epoxy Chip-in-Carrier Integration and Screen-Printed Metalization for Multichannel Microfluidic Lab-on-CMOS Microsystems’, IEEE Trans. Biomed. Circuits Syst., 12(2), pp. 416–425. doi:10.1109/TBCAS.2018.2797063.

Lin, C.-Y. et al. (2022) ‘CMOS Bioelectronics: Current and Future Trends’, in Bioelectronics. Boca Raton: CRC Press, pp. 93–107. doi:10.1201/9781003263265-6.

Mason, A.J. and Wan, H. (2017) ‘Analysis of multi-channel microfluidics for serial dilution in lab-on-CMOS platforms’, in 2017 IEEE 60th International Midwest Symposium on Circuits and Systems (MWSCAS). IEEE, pp. 623–626. doi:10.1109/MWSCAS.2017.8053000.

Ma, W.-T. et al. (2019) ‘The Role of Monocytes and Macrophages in Autoimmune Diseases: A Comprehensive Review’, Frontiers in Immunology, 10. doi:10.3389/fimmu.2019.01140.

Metze, S. et al. (2020) ‘Monitoring online biomass with a capacitance sensor during scale-up of industrially relevant CHO cell culture fed-batch processes in single-use bioreactors’, Bioprocess and Biosystems Engineering, 43(2), pp. 193–205. doi:10.1007/s00449-019-02216-4.

Nabovati, G. et al. (2019) ‘Smart Cell Culture Monitoring and Drug Test Platform Using CMOS Capacitive Sensor Array’, IEEE Transactions on Biomedical Engineering, 66(4), pp. 1094–1104. doi:10.1109/TBME.2018.2866830.

Naresh, Varnakavi. and Lee, N. (2021) ‘A Review on Biosensors and Recent Development of Nanostructured Materials-Enabled Biosensors’, Sensors, 21(4), p. 1109. doi:10.3390/s21041109.

Osouli Tabrizi, H. et al. (2022) ‘Oral Cells-On-Chip: Design, Modeling and Experimental Results’, Bioengineering, 9(5), p. 218. doi:10.3390/bioengineering9050218.

Reinecke, T. et al. (2017) ‘Continuous noninvasive monitoring of cell growth in disposable bioreactors’, Sensors and Actuators B: Chemical, 251, pp. 1009–1017. doi:10.1016/j.snb.2017.05.111.

Reis-Sobreiro, M. et al. (2021) ‘Bringing Macrophages to the Frontline against Cancer: Current Immunotherapies Targeting Macrophages’, Cells, 10(9), p. 2364. doi:10.3390/cells10092364.

Renegar, N., Noyan, U. and Abshire, P. (2022) ‘Deep Neural Network Based Cell Segmentation for Lab-on-CMOS Systems using Realtime Microscopy’, in 2022 IEEE International Symposium on Circuits and Systems (ISCAS). IEEE, pp. 1077–1081. doi:10.1109/ISCAS48785.2022.9937561.

Roussel, T. et al. (2006) ‘3D Microelectrodes for Coulometric Screening in Microfabricated Lab-on-a-Chip Devices’, in 2006 International Conference on Microtechnologies in Medicine and Biology. IEEE, pp. 233–235. doi:10.1109/MMB.2006.251537.

Savitzky, Abraham. and Golay, M.J.E. (1964) ‘Smoothing and Differentiation of Data by Simplified Least Squares Procedures.’, Analytical Chemistry, 36(8), pp. 1627–1639. doi:10.1021/ac60214a047.

Sawan, M., Miled, M.A. and Ghafar-Zadeh, E. (2010) ‘CMOS/microfluidic Lab-on-chip for cells-based diagnostic tools’, in 2010 Annual International Conference of the IEEE Engineering in Medicine and Biology Society (EMBC). IEEE, pp. 5334–5337. doi:10.1109/IEMBS.2010.5626464.

Saylor, J. et al. (2018) ‘Spatial Mapping of Myeloid Cells and Macrophages by Multiplexed Tissue Staining’, Frontiers in Immunology, 9. doi:10.3389/fimmu.2018.02925.

Senevirathna, B. et al. (2016) ‘Lab-on-CMOS capacitance sensor array for real-time cell viability measurements with I2C readout’, in 2016 IEEE International Symposium on Circuits and Systems (ISCAS). IEEE, pp. 2863–2866. doi:10.1109/ISCAS.2016.7539190.

Senevirathna, B., Lu, S., Smela, E., et al. (2019) ‘An Imaging Platform for Real-Time In Vitro Microscopic Imaging for Lab-on-CMOS Systems’, in 2019 IEEE Biomedical Circuits and Systems Conference (BioCAS). IEEE, pp. 1–4. doi:10.1109/BIOCAS.2019.8919023.

Senevirathna, B., Lu, S., Dandin, M., et al. (2019) ‘High resolution monitoring of chemotherapeutic agent potency in cancer cells using a CMOS capacitance biosensor’, Biosensors and Bioelectronics, 142(July), p. 111501. doi:10.1016/j.bios.2019.111501.

Senevirathna, B.P. et al. (2019) ‘Correlation of Capacitance and Microscopy Measurements Using Image Processing for a Lab-on-CMOS Microsystem’, IEEE Transactions on Biomedical Circuits and Systems, 13(6), pp. 1214–1225. doi:10.1109/TBCAS.2019.2926836.

Susloparova, A. et al. (2013) ‘Impedance spectroscopy with field-effect transistor arrays for the analysis of anti-cancer drug action on individual cells’, Biosens. Bioelectron., 40(1), pp. 50–56. doi:10.1016/j.bios.2012.06.006.

Susloparova, A. et al. (2015) ‘Electrical cell-substrate impedance sensing with field-effect transistors is able to unravel cellular adhesion and detachment processes on a single cell level’, Lab Chip, 15(3), pp. 668–679. doi:10.1039/C4LC00593G.

Tarique, A.A. et al. (2015) ‘Phenotypic, Functional, and Plasticity Features of Classical and Alternatively Activated Human Macrophages’, American Journal of Respiratory Cell and Molecular Biology, 53(5), pp. 676–688. doi:10.1165/rcmb.2015-0012OC.

Williams, J.W. et al. (2018) ‘Macrophage Biology, Classification, and Phenotype in Cardiovascular Disease’, Journal of the American College of Cardiology, 72(18), pp. 2166–2180. doi:10.1016/j.jacc.2018.08.2148.

Yin, H., Li, L. and Mason, A.J. (2016) ‘Screen-printed planar metallization for lab-on-CMOS with epoxy carrier’, in 2016 IEEE International Symposium on Circuits and Systems (ISCAS). IEEE, pp. 2887–2890. doi:10.1109/ISCAS.2016.7539196.

